# Overtraining often results in topologically incorrect species trees with maximum likelihood methods

**DOI:** 10.1101/140780

**Authors:** Damien M. de Vienne, Fran Supek, Toni Gabaldon

## Abstract

**Background:** Overtraining occurs when an optimization process is applied for too many steps, leading to a model describing noise in addition to the signal present in the data. This effect may affect typical approaches for species tree reconstruction that use maximum likelihood optimization procedures on a small sample of concatenated genes. In this context, overtraining may result in trees better describing the specific evolutionary history of the sampled genes rather than the sought evolutionary relationships among the species.

**Results:** Using a cross-validation-like approach on real and simulated datasets we showed that overtraining occurs in a significant fraction of cases, leading to species trees that are more distant from a gold-standard reference tree than a previously considered (and rejected) solution in the optimization process. However, we show that the shape of the likelihood curve is informative of the optimal stopping point. As expected, overtraining is aggravated in smaller gene samples and in datasets with increased levels of topological variation among gene trees, but occurs also in controlled, simulated scenarios where a common underlying topology is enforced.

**Conclusions:** Overtraining is frequent in species tree reconstruction and leads to a final tree that is worse in describing the evolutionary relationships of the species under study than an earlier (and rejected) solution encountered during the likelihood optimization process. This result should help develop specific methods for species tree reconstruction in the future, and may improve our understanding of the complexity of tree likelihood landscapes.

## Background

Reconstruction of accurate species trees, representing the evolutionary relationships within a group of species, constitutes a major goal in biology. A species phylogeny is not only the best approximation of a taxonomic classification, but also forms the basis for interpreting how the different species’ traits have evolved, and what evolutionary pressures may have played a role. The advent of molecular sequencing and, more recently, the growing availability of complete genomic sequences have triggered the development of myriad methods for inferring species trees from molecular sequences. The method that is the most commonly used is called “supermatrix”, and consists in concatenating individual genes or protein MSAs into a single large MSA prior to the reconstruction of a unique phylogenetic tree using a Maximum Likelihood or Bayesian tree reconstruction method [reviewed in 1, 2].It is commonly recognized that three main sources of error can occur when using a concatenated set of genes for inferring the history of a group of species. First, the individual genes might not actually have a single evolutionary history [3]. This can be due to a number of evolutionary events impacting different genes in various ways such as, among others, horizontal gene transfers (HGT), incomplete lineage sorting, gene duplications and extinctions [reviewed in 4], but also to artifactual reasons, including sequencing errors, contaminations or wrong orthology detection [reviewed in 5]. Second, the analysis (of one or more of the genes) might be statistically inconsistent due to violation of the applied evolutionary model. Finally, the individual genes being small samples, they may have a non-trivial level of stochastic error. However, to our knowledge, no study so far has evaluated the ability of major phylogenetic tree reconstruction methods to effectively tease apart these sources of error (hereafter called *noise*) from the evolutionary signal of the species under study that one wants to uncover through an optimization process (e.g. search for the Maximum Likelihood (ML) tree). Such a process explores the *likelihood landscape* of trees obtained from a gene or a small number of genes and uses it as a proxy of the real tree-likelihood landscape that one would obtain if using the whole set of genes. The two tree-likelihood landscapes may however differ (Fig. 1A) because of the sources of noise listed above, and it is possible that different methods are differentially affected by this problem.

**Fig. 1.**
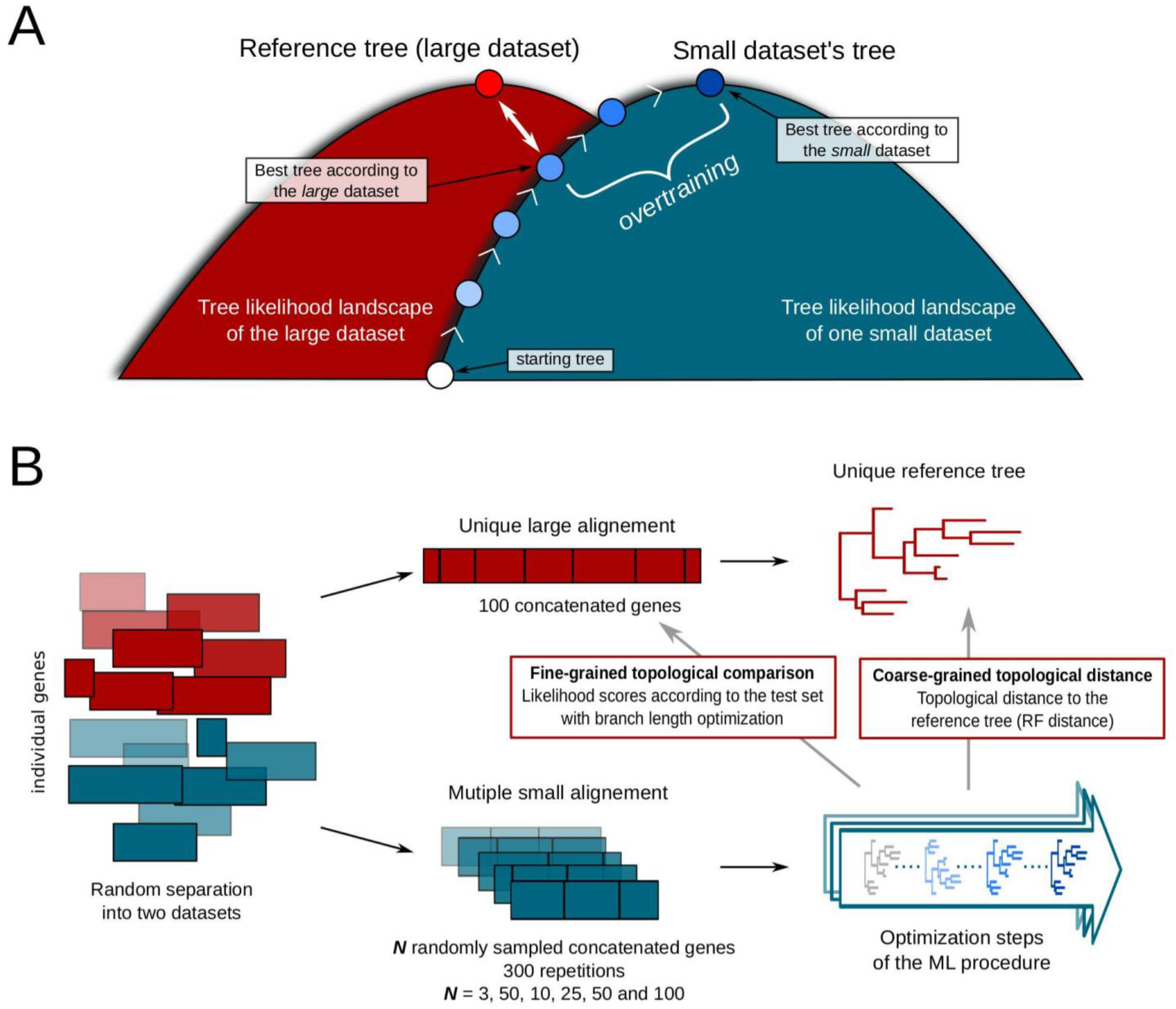
Possible tree likelihood landscapes and description of the method. **A**. Imaginary representation of the difference between the likelihood landscape of a tree reconstructed from a small number of genes (blue hill) and the *real* likelihood landscape of a tree reconstructed from the whole dataset (red hill). The difference in the position of the pick of these two hills may explain the overtraining effect detected in this study. **B**. Schematic description of the method we used to detect overtraining Color code is the same as in A.

A major difficulty faced by all these methods is the need to deal with individual gene histories being not congruent with the underlying species history

Overtraining occurs when an optimization process is trained for too many steps (or epochs), leading to a model describing noise in addition to the signal in the data it was inferred from [6], therefore not generalizing well to new sets of data [reviewed in 7]. In the machine learning field, overtraining was countered by (among others) a measure called “early stopping”, where the training is stopped before the maximum fit to the data was obtained. This procedure is thought to work because the training learns the gross structure of the data first and then the fine structure that is generated by noise [8].

One established statistical technique for evaluating overtraining is the use of a separate data partition for testing the induced model. The cross-validation (CV) technique is an extension of this principle, where the whole data set is repeatedly split into a training and a testing partition prior to the modeling procedure. The measurements of model accuracy on the testing data are accepted as an estimate of accuracy on novel, unseen data.

Overtraining and overfitting are tightly linked concepts that are sometimes considered synonymous. Differences are indeed subtle but are of importance in the present work. First, while *overtraining* leads to a model that overfits, *overfitting* can be obtained without overtraining. Second, overfitting is primarily related to the complexity of a model, while overtraining is primarily related to excessive number of training steps that increase the likelihood of the phylogenetic tree to the sequence alignment, regardless of the complexity of the evolutionary model employed. We do not address here the effect of overly complex models that result in overfitting in the process of phylogenetic reconstruction. This question has been discussed earlier [9] and methods for choosing best-fitting models [i.e. ModelTest, 10] do penalize overly complex models [see 11 for a complete review on model testing]. What we focus on here is the specific impact of overtraining in the context of species tree reconstruction (as opposed to gene tree reconstruction) and how this may affect the quality of the inferred species trees. This question has, to our knowledge, not been studied systematically.

The reconstruction of a species tree from the concatenation of individual gene trees could be prone to overtraining because (i) only a fraction of the data is available (the gene sequences used typically represent a subset of the total number of genes that exist for the species considered) and (ii) noise is present in the data because individual genes are likely to have specific evolutionary histories and biases. The consequence of overtraining are exacerbated as the sample size gets smaller and thus in this case the species tree may not match the phylogenetic signal carried by other gene sequences not used for reconstructing the tree.

We thus set out to investigate the extent of this phenomenon and the effect of size and type of the gene sets thereon. More specifically, we investigated whether overtraining was detectable in the two major ML programs for tree reconstruction, PhyML [12] and RAxML [13] in the context of organismal (species) phylogeny reconstruction.

ML methods for tree reconstruction are optimization procedures that can be described as follows: at each step, the most likely tree from the previous step is used as a starting point for performing topological changes on it, either by Subtree Pruning Regrafting (SPR) or Nearest Neighbor Interchange (NNI). The likelihood of each newly obtained tree (sometimes after a branch length optimization step, depending on the implementation) is then computed and the tree with the highest likelihood is kept for the next step, unless its likelihood is lower than the previously considered tree. A stopping criterion is set for the optimization when the improvement in likelihood between two steps becomes too small.

This optimization results in a final tree whose topology and branch lengths accurately describe the information present in the data (in this case, the aligned gene sequences). Of note, in the context of a species tree reconstruction this might not be the optimal solution, since the aim is to represent with the highest possible accuracy the whole set of data, *i.e*. the complete set of genes, including those not included in the sample used to build the tree. Because an analogy could be made between the optimization procedure of ML tree reconstruction and the training procedures in machine learning algorithms, *overtraining* in this context would refer to a situation in which the number of optimization steps is too high, leading to a final tree representing too specifically the data used to build the tree and conversely losing the ability to generalize and to accurately represent the evolution of the species themselves (Fig. 1A). In other words, overtraining would lead to a final tree (obtained at the end of the optimization procedure) topologically more distant to the species tree than a tree that was met earlier in the optimization.

To investigate this possibility, and assess to what extent overtraining could affect species tree reconstruction, we used a cross-validation approach. We separated each dataset into a large set of concatenated gene alignments used to reconstruct a reference tree, and multiple small sets of concatenated genes not containing the genes present in the large alignment (Fig. 1B). Then, for each small alignment, a tree was reconstructed and the topologies of the intermediate best trees considered during the optimization process were stored (Fig. 1B). By “considered” we refer only to the “current best trees” used as starting point in each of the optimization steps (see above). By comparing the topology of each considered tree to the reference tree (white double-arrow in Fig. 1A) or the likelihood of each encountered tree according to the reference alignment, we can follow the behavior of the ML optimization process and investigate whether overtraining is present. In the absence of overtraining, the topological distance between each considered tree and the reference tree should not increase over the optimization steps; if overtraining occurs, however, the topological distance to the reference tree should decrease at the beginning of the optimization process but then increase when the model becomes too specific to the small alignment. The opposite is expected for likelihood values: if overtraining occurs, the likelihood of encountered trees on the reference alignment should increase at the beginning of the optimization process but then decrease when the model becomes too specific to the small alignment.

The way we define overtraining, *i.e* the presence during optimization of a tree that is closer to the reference tree than the last one, may be seen as too permissive. Indeed, in the hypothetical case where the evolution of the topological distance to the reference tree (or the likelihood of trees on the large alignment) was stochastic, one would expect to often encounter during the optimization a tree that, by chance, is more similar to the reference tree than the last one. To make sure that our observations concern real cases of overtraining and not artefactual ones derived from stochastic tree search paths, we inspected the shape of the likelihood optimization curves. We also used a machine learning approach to demonstrate that some features of the likelihood curves may suggest optimal early stopping points.

## Results

We analyzed three real datasets spanning various evolutionary distances and containing different numbers of taxa: a fungal dataset (246 genes for 21 species [14]), a primate dataset (7018 genes for 10 species, available in PhylomeDB [15]), a cyanobacterial dataset (203 genes for 43 species, [16]), and one simulated dataset derived from the fungal sequences using the program Rose v1.3 [17] following Marcet-Houben *et al*. [18].

### Detection of Overtraining in Tree Reconstruction Methods

For each dataset, we used a maximum likelihood tree reconstruction method (PhyML or RAxML) to reconstruct the maximum likelihood tree and kept track of all the trees encountered during the optimization process. All trees encountered were then compared in terms of topology or likelihood against a reference dataset, either a tree (the reference species tree) or a very large alignment (the reference alignment; see Methods for details).

Regardless of the dataset used, overtraining in terms of tree topology was observable: that is, the final tree returned by the program was often more distant to the reference tree than a tree considered at an earlier point in the optimization process (Fig. 2 and Fig S1). This effect was stronger for smaller sets of concatenated genes and nearly always absent for sets of the same size as the large alignment (Fig. 3). The latter result confirmed that the reference tree was reconstructed from a sufficiently large number of genes. For the simulated dataset, where all genes followed the same evolutionary history, overtraining still occurred, albeit to a limited extent. This indicates that overtraining cannot be completely explained by differences in the underlying gene evolutionary histories, even if increasing the proportion of alternative topologies under simulated controlled conditions clearly leads to an increase in the proportion of cases where overtraining is detected (Fig. S2 and Fig. S3). In addition, the two major reconstruction implementations produced similar patterns of overtraining when applied to the same dataset (Fig. 2), revealing that the problem we point out is not software-specific but rather more generally present in the ML methodology when the goal is to reconstruct a species tree from a small sample of genes.

**Fig. 2.**
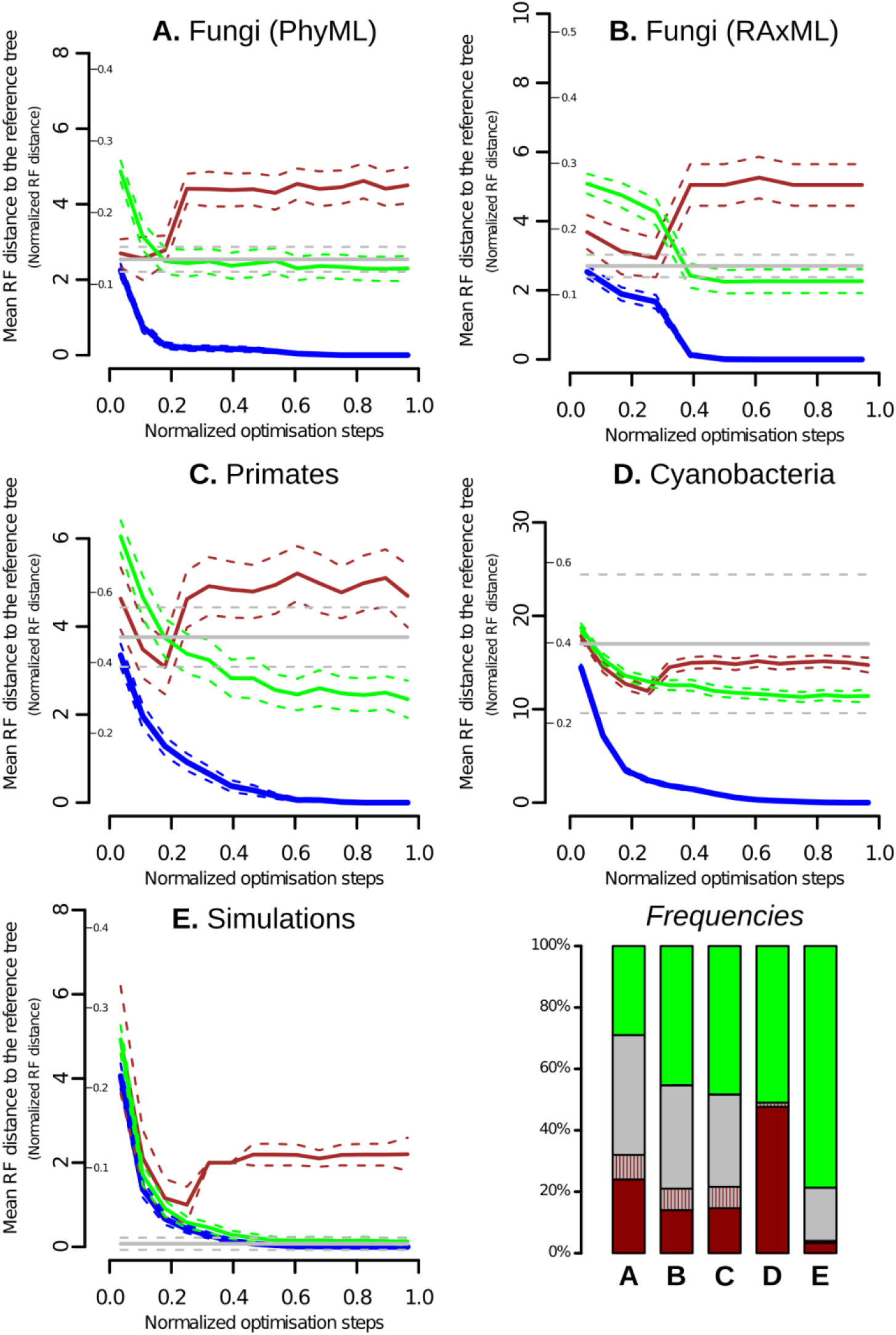
Overtraining detected using RF distances. Illustration of the overtraining effect detected in this study using the RF distance (*y*-axis) between each considered tree (*x*-axis) and the reference tree. The 300 independent runs performed for each dataset are separated into 3 sub-categories: cases where overtraining is detected (red line), cases where the distance between each considered tree and the reference tree is stable along the optimization (grey lines) and cases where an improvement in topology is detected (green lines). The blue lines represent topological distances between each considered tree and the last tree obtained. All examples are for training alignments of size five (5 concatenated genes, see Fig. 1b). The *x*-axis is normalized so that the first tree is assigned the value 0 and the final tree the value 1. Moreover, for each independent run, the x-axis is stretched so that the position of the best tree considered has always the same *x* value. This value is the mean position of the best considered tree over all runs. The mean RF distance to the reference tree is then computed after separating the x-axis into bins of size 15 for all datasets (except B. for which the number of bins is 10). The bar plot on the bottom right corner represents the frequency of each scenario for each dataset tested. The portion with red dashes depicts the fraction of the runs where the topological distance to the reference tree is stable but where overtraining is detected if looking at the likelihood of the trees. On panels A to E, the small graduation on the y-axis shows RF distances normalized by the number of nodes in the trees.

**Fig. 3.**
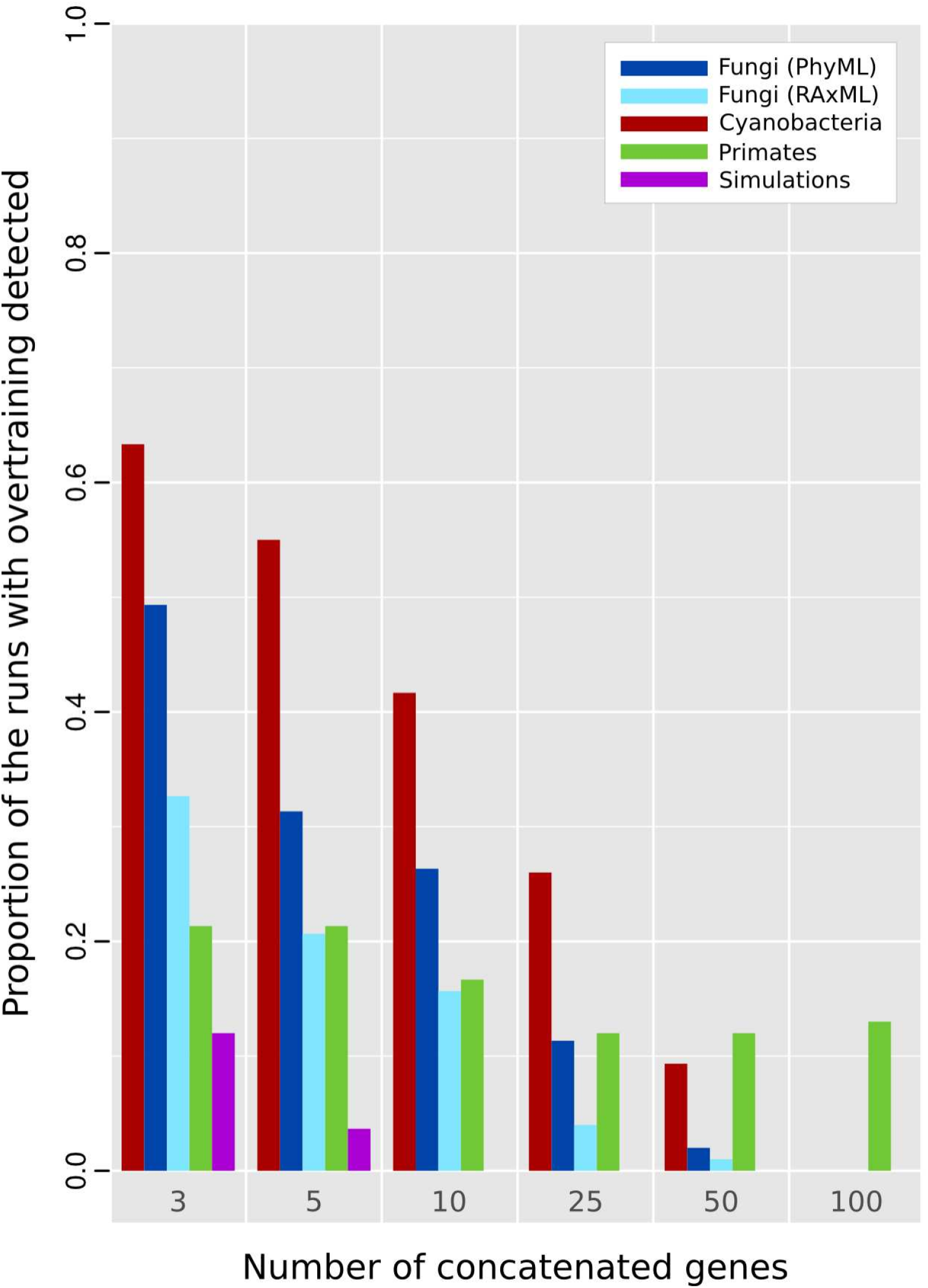
Total frequency of overtraining. The proportion of runs where overtraining is detected either with RF distance or with Likelihood measures is shown, for each size of the learning alignments and each dataset used in this study.

### Refined Measure of Topological Overtraining Using Likelihood Values

The above measures can be considered a conservative estimate of the degree of overtraining because the Robinson-Foulds (RF) measure [19] used to compute the topological distances lacks precision. Indeed, trees having different topologies may have the same RF distance to the reference tree, despite one of them being a better fit in terms of likelihood. In order to examine these cases in more detail, we measured the likelihood of each tree topology considered during the optimization process based on the large concatenated alignment (Fig. 1A). In order to specifically measure topology only we initially set all branch lengths to 1 and allowed them to vary freely during the optimization process. Each optimization run then falls into different categories, depending whether it shows topological overtraining based on RF distance and/or topological overtraining based on likelihood scores (see Fig. S4 for possible scenarios). As the RF metric is a more coarse-grained measure of topological distance, runs where topological overtraining was detected using RF distances were also showing topological overtraining in terms of likelihood (Fig. 4 and Fig. S5). Additionally, some runs that did not show overtraining in terms of RF did so according to likelihood distances (Fig. 4 and Fig. S5).

**Fig. 4.**
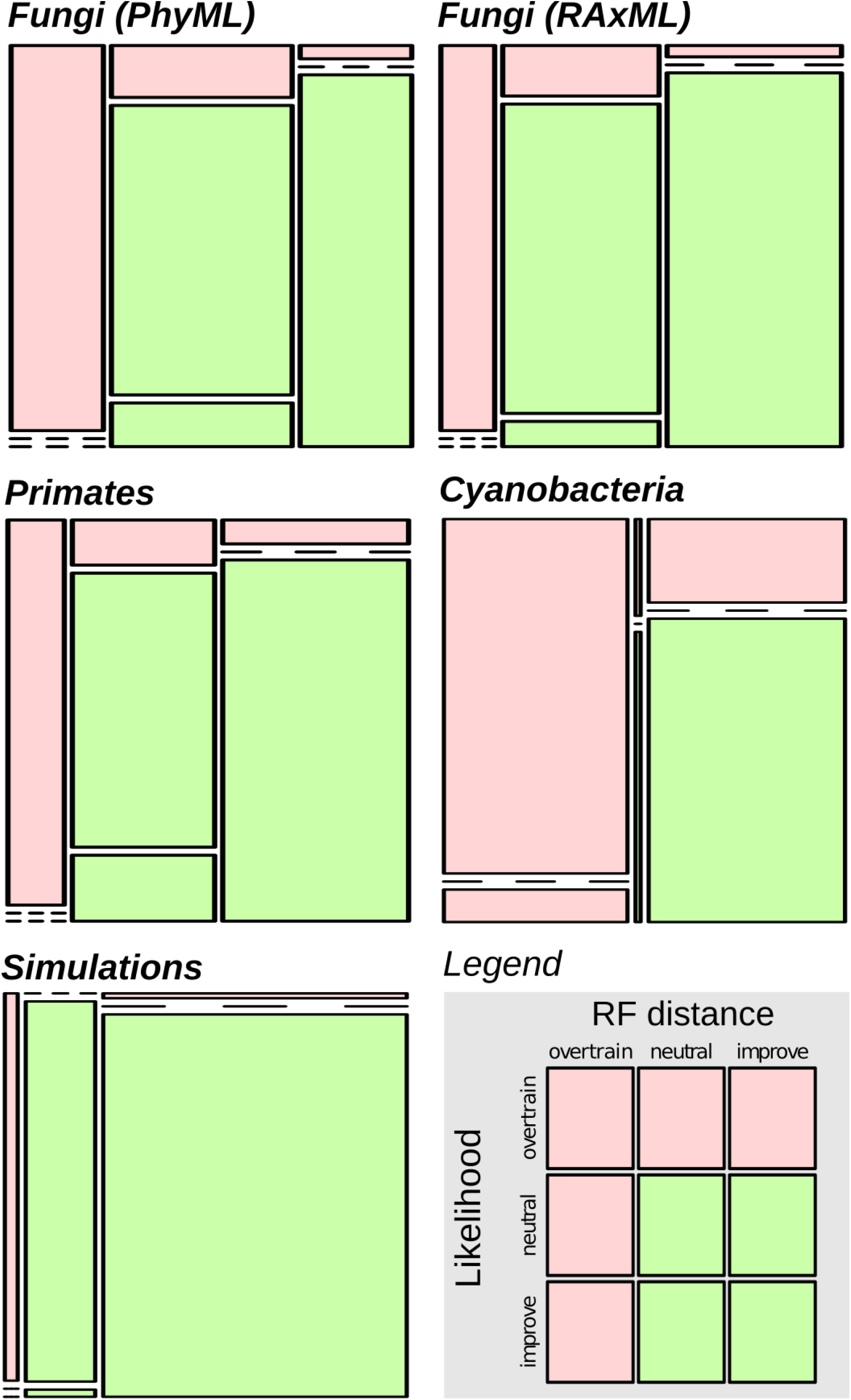
Overtraining detected using RF distances and likelihood. Mosaicplot showing the proportion of runs where topological overtraining occurs for the different datasets and for the two detection methods applied: RF distance (columns) or likelihood (rows). All examples are for training alignments of size 5. A detailed legend is given in the bottom right panel.

### Overtraining and Node Support

It is a common procedure in phylogenetics to evaluate the quality of a tree by measuring the stability of each node to data resampling. To investigate whether the nodes that are incorrect in the final tree due to overtraining are distinguishable and could thus be corrected or removed *a posteriori*, we inspected bootstrap and approximate Likelihood Ratio Test (aLRT) supports in the fungal dataset. As expected, the incorrect nodes (with respect to the reference tree) had much lower support than the correct ones but among them, nodes introduced by overtraining (see Material and Methods) were practically indistinguishable by their bootstrap support values from the other incorrect nodes (Fig. S6A), and also by their aLRT support (Fig. S6B). Therefore, node support metrics cannot be used to single out the overtraining-induced errors in tree topology. Note moreover that, in this context, the aLRT method was a worse “distinguisher” of incorrect nodes than the classical bootstrap approach, despite having been shown to correlate with the bootstrap [20]. Furthermore, by following the aLRT support values of trees along the optimization process, we could show that the number of highly supported nodes increases (Fig. S7). The fact that the considered trees are progressively more resolved during the optimization process could be interpreted as an increase in the complexity, which is consistent with overtraining being a cause for the observed drop in model accuracy as training progresses past the optimum.

### Overtraining is not simply a stochastic effect

We tested whether the overtraining effect that we detected could be simply the result of stochastic effects during the tree optimization process.

There are two possible effects that could create overtraining-like signal. First, by chance alone, in the path from the initial tree to the final ML tree, some search runs will show trees passing closer to the reference tree and these runs will be erroneously identified as overtraining by our approach. If this is true, the effect should be independent of the reference tree topology. That is, the proportion of detected cases with such apparent overtraining should be constant regardless of the reference tree topology chosen. To test this, for the fungal dataset and a small size of alignment (5 genes), we generated 100 random trees and evaluated each time the number of runs where overtraining was detected (Fig S8). We observed that chance alone could account for only about 1% of the runs considered as overtraining (mean = 0.012, s.d. = 0.03) while the observed value was 24% (72 runs out of 300).

Second, if the evolution of the RF distances to the reference tree (or the likelihood values against the large alignment) during the optimization was purely stochastic, that is bumping up and down in a random fashion, overtraining would thus also be obtained in a non-negligible fraction of cases. To ensure that this is not what is happening, we used a machine learning strategy to show that the evolution of likelihood values on the small alignment was actually a very good predictor of the position of the optimal species tree. We calculated 47 features (see Methods and Table S1) that described the shape of the likelihood curves in each given training step (illustrated in Fig S1), and used these to train a Random Forest classifier to detect the optimal stopping point in each run. The cross-validation AUC values that describe how accurate the classifier is at discriminating between best-tree (closest to reference) and all other trees encountered during training were almost all superior to 0.9 (Table 1). This rules out the possibility of stochasticity-induced (and thus erroneous) overtraining effect. To ensure that what we are learning is not simply from the runs where no overtraining is detected, so that what we learn would simply be that the last tree is the best, we redid all learnings with only runs where overtraining was present. The results were similar, with even higher AUC values in most cases (Table 1).

**Table 1.**
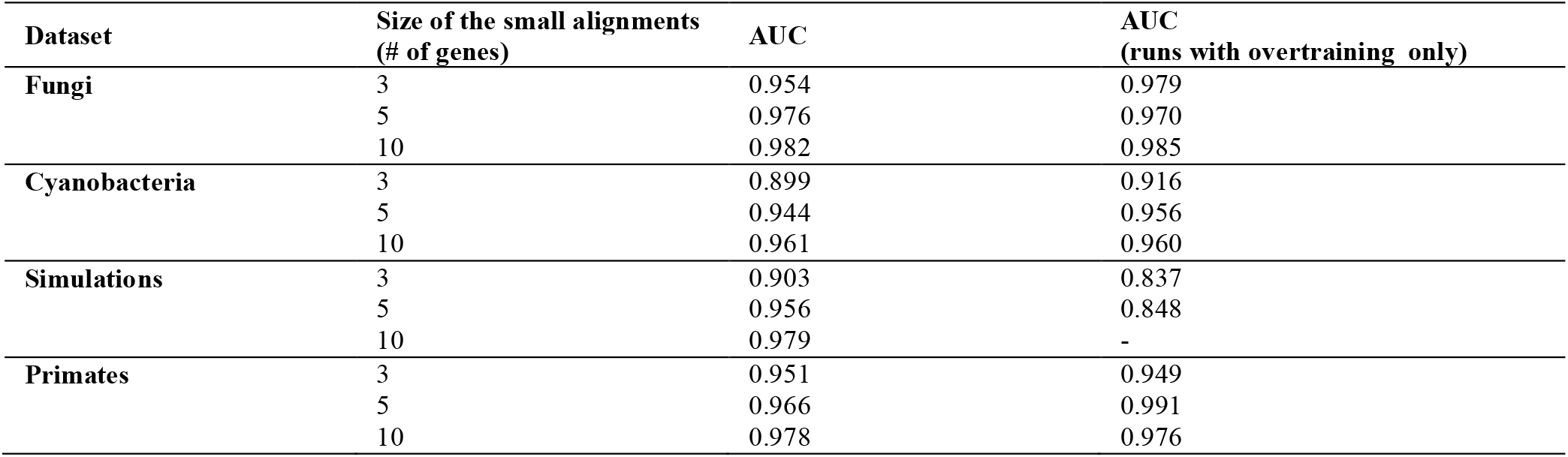
AUC scores of the Random Forest Classifier for predicting the best tree from the likelihood values on the small alignments.

| Dataset | Size of the small alignments<br>(# of genes) | AUC | AUC<br>(runs with overtraining only) |
| --- | --- | --- | --- |
| Fungi | 3 | 0.954 | 0.979 |
|  | 5 | 0.976 | 0.970 |
|  | 10 | 0.982 | 0.985 |
| Cyanobacteria | 3 | 0.899 | 0.916 |
|  | 5 | 0.944 | 0.956 |
|  | 10 | 0.961 | 0.960 |
| Simulations | 3 | 0.903 | 0.837 |
|  | 5 | 0.956 | 0.848 |
|  | 10 | 0.979 | - |
| Primates | 3 | 0.951 | 0.949 |
|  | 5 | 0.966 | 0.991 |
|  | 10 | 0.978 | 0.976 |

### Bayesian approaches are less prone to overtraining but do not give more accurate species trees

We investigated whether Bayesian approaches [here MrBayes, 21, 22] were less prone to overtraining than ML, as expected for an approach that samples over a likelihood landscape. This was indeed the case, and the proportion of runs with overtraining was approximately 5% with MrBayes, as compared to 24 to 36% for PhyML on the same datasets (Fig. S9 A). However, this does not necessarily imply that Bayesian approaches are better at reconstructing species trees than ML approaches. Indeed, when comparing the topological distances between the MrBayes final (consensus) trees and the reference tree in the one hand and the topological distances between the final ML trees and the reference tree in the other hand, we observed that for most runs, the trees were identical and that when they differed, MrBayes or PhyML trees were similarly likely to be closer to the reference tree (Fig. S9 B).

### Exhaustive search of the most likely tree and visualization of tree likelihood landscapes

To ensure that overtraining was not attributable to a particular implementation of the ML heuristics, we performed an exhaustive search for the most likely tree for a fungal dataset of 9 species and compared to the ML tree obtained by RAxML after a normal run. We focused on 25 alignments of 5 genes where overtraining was present. For all the alignments, the final trees were the same in the normal and exhaustive searches and the final tree was always the most likely in the exhaustive search. This test also allowed us to visualize the entire tree likelihood landscape for each one of these 25 runs, by performing a multidimensional scaling (MDS) on the pairwise distance matrix of the best trees encountered (see Material and Methods). Such a landscape is given in Figure 5. We observe different “islands” of higher likelihood and see that the most likely tree given the 5-gene alignment (white arrow) is at a different peak than the reference tree (black arrow). When looking at the 25 landscapes (Fig. S10), we observe that even though the species tree is never the one with highest likelihood (because we chose runs with overtraining), it is always at an apparent local optimum. Moreover, when looking at the mean rank of all trees for the 25 analyses (Figure S11), the reference tree is the one with the smallest mean rank and the smallest standard deviation (mean = 2.64; sd = 1.11). The best ML trees of each run display higher mean ranks.

**Fig. 5.**
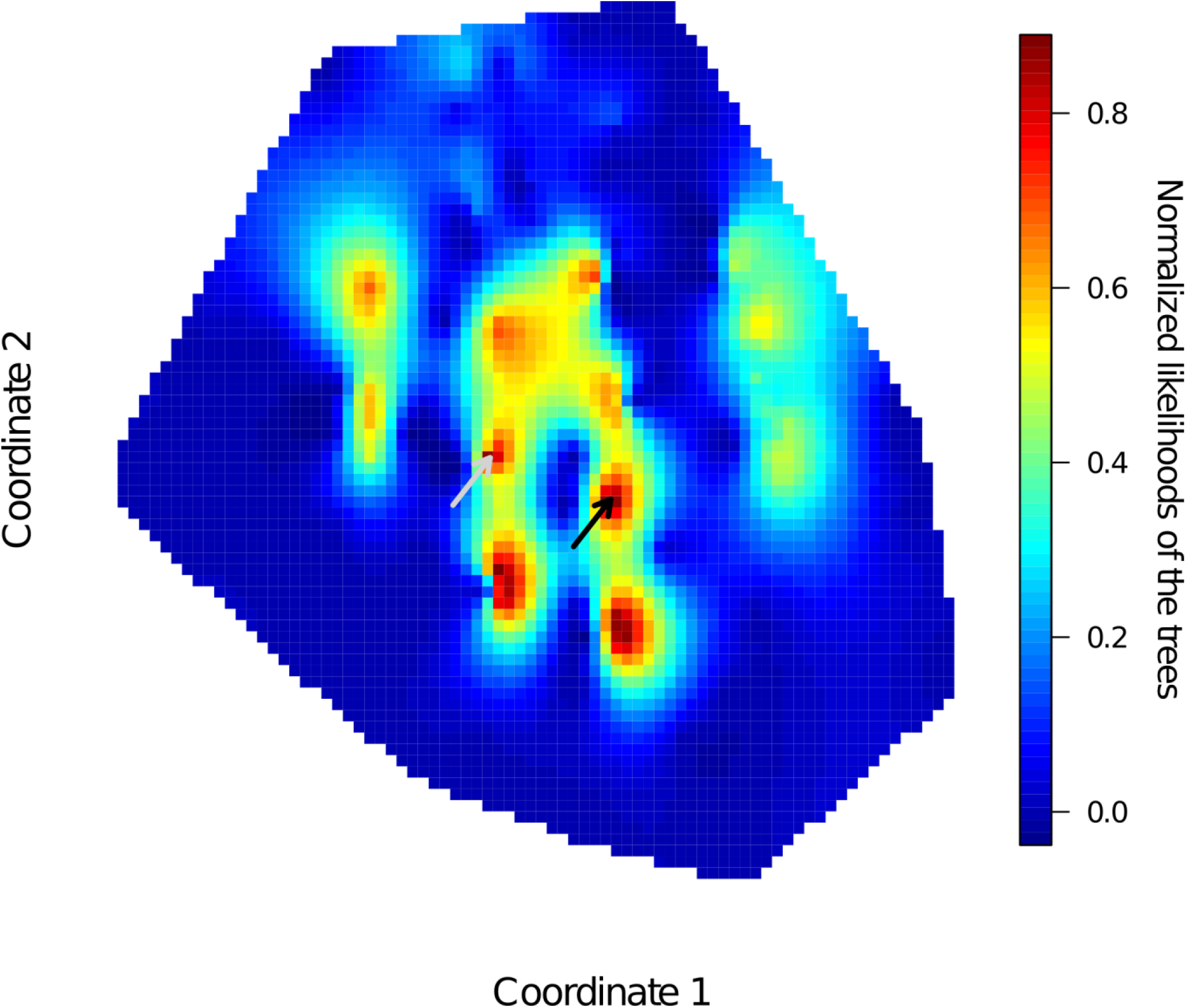
Representation of the tree likelihood landscape for an alignment of 5 concatenated genes for 9 fungal species. The most likely tree (white arrow) is distinct from the reference species tree (black arrow). Hot colors depict zones of higher likelihoods. The scale on the right panel represents normalized likelihood values.

Therefore, with this dataset, the reference tree is never the best (judged by the fit on the 5-gene alignment), but always has a high likelihood: that is, it seems to be robust to gene sampling effect. This observation may provide ideas for the development of future solutions to the overtraining problem.

## Discussion and Conclusion

Using a simple approach wherein we reconstructed phylogenetic trees from progressively smaller samples of genes and compared them to a reference tree reconstructed from a large gene set, we have shown that overtraining is common in methods based on Maximum. Consequently, a tree reconstructed using a small number of concatenated genes, as is done on a regular basis, often results in species trees more distant to the reference tree than what could be obtained if overtraining was curtailed in some manner. Of note, this does not imply that the optimization process did not improve the tree at all, but rather indicates that the best considered solution was often missed due to additional rounds of unnecessary iterations, in addition to the wasted computational time.

In our view, future versions of the major ML reconstruction methods would benefit by explicitly taking overtraining into account when the goal is to recover the organismal phylogeny and not the specific evolutionary history of a small set of genes. Possible implementations that would alleviate this problem include finding early stopping criteria, or constraining the complexity of the evolutionary model prior to (or even during) the optimization.

Our results suggest that a large fraction of the species trees present in the literature, especially those based on small sets of genes, have overtraining-related topological errors. This may explain, at least in part, the topological incongruities between species trees published by different research groups for the same set of species: overtraining can contribute to different sets of concatenated genes resulting in different species trees.

## Material and Methods

### Datasets

#### Fungal dataset

The fungal dataset [14] was obtained from the authors as a set of already aligned and trimmed multiple sequence alignments (See [14] for the methods employed). Species included in this dataset (Table S2) span a wide range of the Fungal tree of life.

#### Primate dataset

This dataset was retrieved from Phylome-DB database [15], Phylome-DB phylome accession number: 98. It was obtained using the protocol detailed in [23]. Only one-to-one orthologous sequences present in all 10 primate species considered (Table S2) were kept, resulting in 7,018 genes. Each set of orthologous proteins was aligned using MUSCLE version 3.7 [24] with default parameters and trimmed using trimAl v1.4.rev7 [25] with the gappyout option (automatic detection of blocks to remove based on the distribution of gaps in the alignment).

#### Cyanobacterial dataset

This dataset described in [16] was retrieved from Phylome-DB database [15], Phylome-DB phylome accession number: 30. It was constructed using all available cyanobacterial complete genomes (listed in Table S2). The same method was used for the detection of orthologous sequences as for the Primate dataset (see [23] for a description of this method).

#### Simulated dataset

We used the program ROSE v.1.3 [17] to generate multiple alignment for 21 species using a known fungal phylogeny [18] (Fig. S1, left tree). Sequences, trees and mutation probability of each amino-acid in the input sequence were taken from [18]. ROSE was run with default parameters. To obtain a realistic set of alignments, we tuned the branch lengths of the input tree so that the simulated sequences had the same distribution of similarity scores, after alignment and trimming, as the alignments present in the fungal phylome of PhylomeDB database [15] from which the input sequences were taken. This was achieved by multiplying all branch lengths by 4.

### Tree Reconstruction and Overtraining Evaluation

The principle of the method used in this study to evaluate overtraining is presented in Figure 1. All the analyses were performed using PhyML version 3.0 [12]. For the Fungal dataset, RAxML version 7.2.8 [13] was used in addition to PhyML.

#### Construction of the reference tree from the large alignment

The concatenated alignment of 100 genes (or 200 for the primate dataset) was used to reconstruct a reference tree using either PhyML [12] or RAxML [13]. The PhyML parameters were -m LG -b 0 -f e -v e -a e -o tlr -c 4 and the RAxML parameters: -*m PROTGAMMAIWAGF*.

#### Construction of the trees from the small alignments

Exactly the same parameters were used in this case but all the trees considered during the optimisation were stored, using the parameter -*print_trace* (PhyML) or -j (RAxML). Note that we call “trees considered” only the “current best trees”, that are trees whose likelihood is the best at each step and that are thus used as starting trees for the next step.

#### Estimation of overtraining of trees based on topological distance

The topological distances between each tree considered (*T_search_*=[*T*_1_, *T*_2_, *T*_3_, …., *T_n_*]) and the reference tree (*T_ref_, obtained from the large concatenated dataset)* was computed using the Robinson-Foulds (RF) distance [19] as implemented in the R package “ape” [26]. We considered as “*overtraining*” cases where there exists at least one *i* in [1*(n*−1)] so that

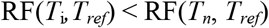

*Estimation of overtraining of trees based on likelihood*.—Two topologically different trees can have the same topological distance to the reference tree and be therefore indistinguishable. To have a finer measure of topological overtraining, we computed the Log-likelihood (-LogLK) of each considered tree (each topology) according to the large alignment after having set all branch lengths to 1. We only fixed the topology and let the branch length and model parameters be optimized by the software used. PhyML was executed with the following parameters: -*m LG -b 0 -f e -v e -a e -o lr -c 4* and RAxML with -*m PROTGAMMAIWAGF -f e*, using the large alignment as the input and each considered tree as the user-defined tree topology. If we call *T_search_*=[*T*_1_, *T*_2_, *T*_3_, …., *T_n_*] the trees considered during the optimization and MSArefthereference multiple alignment, then we considered as overtraining cases where there exists at least one *i* in [1,(*n*−1)] so that

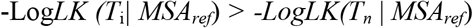

### Bootstrap and aLRT Analyses

To investigate whether an incorrect node caused by the overtraining could be distinguished from an incorrect node due to other causes, for each run where overtraining (in terms of RF distance) was detected, we computed the bootstrap support and the aLRT support of the nodes in the final tree – the tree normally obtained by the user. This analysis was performed for the Fungal dataset using PhyML and 100 bootstrap replicates for the Bootstrap analysis and the aLRT non-parametric branch support based on a Shimodaira-Hasegawa-like procedure for the approximate Likelihood Ratio Test analysis, as implemented in PhyML (option -*b* = −4).

To distinguish between an incorrect node due to overtraining and an incorrect node due to alternative causes, the nodes in the best considered tree were compared to the reference tree and to the final tree. If a node was correct in the best considered tree (according to the reference tree) and incorrect in the final tree (according to the reference tree again), we considered that overtraining was responsible for the creation of this incorrect node.

### Link Between the Proportion of Alternative Individual Gene Histories and Overtraining

To test whether the proportion of genes following an alternative evolutionary story that the reference one had an impact on the amount of overtraining, we simulated sequences using ROSE with the same input parameters as for the simulated dataset described above but with two different input trees: the “reference topology” and the “alternative topology” (Fig. S1). The alternative topology was obtained from the reference topology by moving a clade of 4 species to another position in the tree (red dots in Fig. S1). We only considered the training set where 10 genes were concatenated and we varied the proportion of alternative topologies (0%, 10%, 20%, 30% and 40%) included in the training set. 300 repetitions were performed each time and the proportion of runs where overtraining occurred (detected with RF distances) was estimated.

### Classification of Overtraining Scenarios

We called *overtraining* those cases were at least one of the trees considered during the optimization process was better than the final tree. We called *improvement* those cases where the last tree was the best one and where at least one of the previous trees was worse. We called *neutral* those cases where all trees were equally good (or bad) all along the optimization process. The qualifiers “better” or “worse” can be seen in terms of RF distance or in terms of likelihood of the tree given the large alignment. Figure S4 gives a schematic representation of all these scenarios. Some of them, indicated by (*) are not realistic: when the likelihood of the trees do not change along the optimization process while the topological distance to the reference tree changes. Indeed, it is largely unlikely that two trees with different topologies have exactly the same likelihood according to a given alignment.

### Test for potential random effects leading to overtraining detection

To ensure that the overtraining effect we detect is not random, *i.e*. is not due to paths from NJ tree (or MP trees in the case of RAxML) to ML trees passing by chance close to the reference tree, we evaluated the proportion of runs where overtraining was detected when replacing the reference tree by random trees. This was done 100 times, using the the R package ape (<u>29</u>) to generate the random trees. We used the paths obtained for the 5 concatenated genes from the fungal dataset for this test. The obtained null distribution was then compared to the observed value.

Another source of artifact could be if the evolution of Robinson-Fouldsdistances against the reference tree (or Maximum Likelihood values against the large alignment) during optimization was stochastic. In this case, we may detect cases of overtraining quite often just by chance.

We used a Random Forest classifier on the curves describing the evolution of likelihood values on the small alignments to try to predict where the best stopping point was. In particular, for each point in the optimization, we computed 47 features: 43 that described the shape of the lk curve in the vicinity of that specific point, and 4 more features that described the curve globally (Table S1). We used the *randomForestSRC* package in R [27], with *ntree*=200, *na.action*=“na.random”, *importance*=”permute” and all other parameters left at default values. To find the AUC score, we used the out-of-bag predictions (equivalent to leave-one-out cross-validation) stored in *predicted.oob*, and processed them using the *ROCR* package [28]. Features are presented in Table S1.

We wondered whether the good predictions we obtained could be due to simply detecting those runs where there was no overtraining in contrast to the runs where this happens, while not necessarily being able to distinguish the good stopping points from the bad ones in the overfitting runs. To test, we removed non-overtraining runs and re-did the Random Forest learning and testing. The results obtained were similar, proving that even when overtraining is present, the likelihood values on the small alignment allow predicting the best tree according to the large alignment (or, equivalently, the tree closest to the reference tree).

### Evaluation of overtraining with Bayesian approach

For comparison with ML methods, we used MrBayes v3.2.2 [21, 22] on the fungal dataset for small alignments of size 3 and 5. For each size and each of the 300 repetitions, MrBayes parameters were chosen as follows: 2 runs, one single chain, 1.000.000 generations, and sampling every 500 generations. The model used was the same as for the PhyML tests: fixed LG substitution matrix (added in the source code by ourselves), gamma-distributed rate variation across sites and a proportion of invariable sites (rates=invgamma). Five runs did not reach convergence after 1.000.000 generations (average standard deviation of the split frequencies higher than 0.01) and were discarded.

For comparison with the ML approach, the consensus species tree was computed after keeping only the first 5%, 10%, 15%…., 95%, 100% of the trees in the two runs, burning always the first 25%. This led to 20 trees for each run for which we could detect overtraining based on RF by comparison with the reference tree.

We also compared the final tree obtained with MrBayes to the final tree obtained with PhyML in terms of their RF distance to the reference tree, to see whether one of the two methods was naturally better at capturing the evolutionary history of the species than the other.

### Exhaustive search of the most likely tree for 9 species

We modified the 300 alignments for the 5 concatenated fungal genes; in order to keep only 9 species (Species in bold in Table S2). RAxML was then used to detect overtraining with these reduced alignments, using the same approach as before. Twenty-five runs for which overtraining was detected were analyzed further. We generated, using Paup v4.0 beta [29], all possible unrooted tree topologies containing 9 species (135135 trees) and computed for each one its likelihood according to each of the 25 alignments. We then compared the ML tree obtained with RAxML for each run to the real best tree in the exhaustive search.

We also used these data to visualize “real” likelihood landscapes. To do so, we selected the 100 best trees for each run (leading to 376 unique topologies) and computed all against all RF distances between these trees using Paup v4.0 beta [29]. This led to a distance matrix on which we performed multidimensional scaling (MDS) to obtain a two-dimensional representation of the tree landscape. We finally applied a kriging method on the MDS data as implemented in the R package *fields* [30] in order to better visualize the landscape (as a surface and not as isolated points). This is what is represented in Figure 6 and Figure S10.

Finally, we computed for each of the 135.135 topologies their rank in the sorted list of likelihoods for each of the 25 alignments. The mean and standard deviation of these ranks were then calculated and sorted (Figure S11).

### Statistical difference between best and last ML trees in cases of overtraining

We wanted to assess whether the last ML tree in cases of overtraining was significantly different from the best tree encountered during the optimization, with respect to the large alignment. For a subset of 50 overtraining runs (fungal dataset, 5-genes alignments), we selected the best ML tree and the last ML tree. We then used PhyML to compute sitewise likelihood of the large alignment according to each tree, letting the rate and branch length to be optimized, and finally input this into CONSEL v1.20 [31]. CONSEL allows comparing alternative topologies by performing the approximately unbiased (AU) test and computing the associated p-value. At a significance level 0.05, we observed that for 60% of the runs (30 out of 50), the final ML could be excluded. This shows that overtraining is not simply a “waste” of optimization time. It is also producing trees that are significantly worse than previously encountered ones in a large fraction of cases.

## Availability of data and materials

The data reported in this paper are available at the following address: ftp://ftp.cgenomics.org/overfit/.

## Competing interests

The authors declare that they have no competing interests

## Funding

TG research group is funded in part by the Spanish Ministry of Economy and Competitiveness (BIO2012-37161), a Grant from the Qatar National Research Fund grant (NPRP 5-298-3-086), and a grant from the European Research Council under the European Union′s Seventh Framework Programme (FP/2007-2013) / ERC (Grant Agreement n. ERC-2012-StG-310325). FS was co-funded by Marie Curie actions.

## Authors′ contributions

DMDV, FS and TG conceived the research, DMDV performed the analyses, DMDV, FS and TG analyzed the results and wrote the manuscript.

## Consent for publication

Not applicable.

## Ethics approval and consent to participate

Not applicable.

## Acknowledgements

We would like to thank A. Stamatakis and O. Gascuel for comments on an earlier version of this manuscript, and Dragan Gamberger, Tomislav Šmuc, Franck Picard and Laurent Jacob for discussion on machine learning and overtraining in the phylogenetic context.

